# Modulation of Receptor-like Trans-Membrane Kinase 1 nuclear localisation by DA1 peptidases in Arabidopsis

**DOI:** 10.1101/2022.04.01.486729

**Authors:** Benguo Gu, Hui Dong, Caroline Smith, Michael W. Bevan

## Abstract

The activities of proteins can depend on their cleavage, which influences their cellular location, stability and functions. The cleavage of intracellular domains of Receptor -Like Kinases (RLK) has an important functional role in the transduction of signals from the cell surface to the nucleus in many organisms. However, the peptidases that catalyse protein cleavage during signal transduction remain poorly understood despite their crucial roles in diverse signalling processes. Here we report in the flowering plant *Arabidopsis thaliana* that members of the DA1 family of ubiquitin-regulated Zn metallopeptidases cleave the cytoplasmic kinase domain of TransMembrane Kinase 1 (TMK1), releasing it for nuclear localisation where it represses auxin-responsive cell growth during apical hook formation by phosphorylation and stabilisation of the transcriptional repressors IAA32 and IAA34. Mutations in *DA1* family members exhibited reduced apical hook formation, and DA1 family mediated cleavage of TMK1 was promoted by auxin treatment. Expression of the DA1 family - generated intracellular kinase domain of TMK1 by an auxin-responsive promoter fully restored apical hook formation in a *tmk1* mutant, establishing the function of DA1 family peptidase activities in TMK1-mediated differential cell growth and apical hook formation. DA1 family peptidase activity therefore modulates TMK1 kinase activity between a membrane location where it stimulates acid cell growth and initiates an auxin-dependent kinase cascade controlling cell proliferation in lateral roots and a nuclear localisation where it represses auxin mediated gene expression and growth.

## Introduction

Protease-mediated cleavage is essential for the functions of many diverse proteins, including their trafficking to cellular compartments, protein homeostasis, and the relocation of protein domains. In many signal transduction pathways involving the large family of membrane-localised Receptor Tyrosine Kinases (RTKs), proteolytic cleavage is essential for their signalling functions (1). Upon activation by extracellular signals, ectodomains can be shed to the extracellular region by a variety of proteases including alpha-secretases (2). This can be followed by further proteolytic cleavage by the membrane-located gamma-secretase complex to form intracellular domains (ICD) that are released into the cell to perform diverse regulatory functions (3). Although flowering plants encode gamma-secretase subunits (4, 5) and Rhomboid intramembrane serine proteases that have highly conserved functions in proteolytic processing of transmembrane regions of specific proteins (6), their substrates in plants are unknown.

Plants encode multiple membrane-localised Receptor-Like Kinases (RLKs) that detect stimuli from pathogens, symbionts (7) and endogenous signals (8, 9), transducing these into cellular responses. The intracellular kinase domains of membrane-localised RLKs generally initiate signalling from the plasma membrane by phosphorylation of membrane -associated proteins and by kinase cascades from the plasma membrane, for example in brassinosteroid (BR) signalling (10). RLK kinase domains can also be processed into ICDs that function elsewhere in the cell. For example, the rice XA21 receptor kinase confers resistance to bacterial pathogens and treatment of plants with an XA21 ligand led to the release of a C-terminal kinase-containing ICD that localised to the nucleus where it interacted with a transcription factor (11). The intracellular domain of the RLK BAK1 is cleaved by a conserved Ca^2+^-dependent protease activity close to the transmembrane domain to release an ICD (12). An amino acid substitution that abrogated BAK1 cleavage reduced pathogen- and BR-mediated responses and influenced the plasma membrane localisation of BAK1, indicating a role for proteolytic cleavage in BAK1 function. The identity of proteases involved in XA21 and BAK1 cleavage has not yet been determined. Over-expression of the secreted carboxypeptidase BRS1 suppressed weak alleles of the RLK BR receptor BRI1 and this suppression was lost by mutations in the conserved peptidase active site (13). This suggested that peptidase activities were required for early stages of BR signalling, although no proteolytic processing of BRI1 has been reported (14).

There are two relatively well-described cases of proteolytic processing of membrane signalling proteins in plant hormone signalling pathways. EIN2 is a transmembrane protein located in the endoplasmic reticulum (ER) that is essential for transducing signals from the ethylene receptor. It has a C-terminal domain that is proteolytically released from the ER in response to ethylene and functions to modulate the activities of transcription factors EIN3 and EIL1 by direct interaction in the nucleus and in cytosolic P-bodies to enhance the degradation of *EBF1/2* mRNA, which function to modulate EIN3 and EIL1 protein stability (15–18). Subsequent studies identified multiple C terminal domain EIN2 protein fragments in the absence of ethylene treatment, and full-length EIN2 was also observed in the cytosol and nucleus, revealing a more nuanced and complex role of EIN2 proteolysis in ethylene signalling (19). The RLK Transmembrane Kinase 1 (TMK1) phosphorylates plasma membrane H+ -ATPases in response to auxin and activates cell wall acidification, cell expansion and growth ((20–22). TMK1 (redundantly with its family member TMK4) also phosphorylates MKK4/5 in response to auxin and initiates a kinase cascade that regulates cell divisions during lateral root formation (23). TMK1 kinase activity is inhibited by membrane-associated kinase regulator 2 (MAKR2) to influence root gravitropism and auxin antagonises this inhibition by dissociating MAKR2 from the membrane (24). In addition to these membrane-localised kinase activities of TMK1 that influence cell growth, the cytosolic kinase domain of TMK1 is also cleaved by an auxin-dependent mechanism and translocates to the nucleus, where its kinase activity phosphorylates and stabilises IAA32 and IAA34 transcriptional repressors that are part of the TIR/AFB signalling pathway, leading to reduced cell growth and apical hook formation (25). The identity of the protease(s) involved in switching TMK1 kinase activity from a membrane-localised promoter of cell growth to a nuclear-localised repressor of TIR/AFB transcriptional responses is currently not known, despite a pivotal role in modulating auxin dependent growth processes mediated by TMK1.

The DA1 family of peptidases is highly conserved in plants and three family members, DA1, DAR1 and DAR2 function redundantly to limit cell proliferation during organ growth by cleaving a variety of growth promoting proteins that are subsequently targeted for degradation (26, 27). The peptidase activity of DA1 is activated by monoubiquitylation (27), reduced by de-ubiquitylation (28), and inhibited by BRI1-mediated phosphorylation (29), indicating a potential regulatory role for DA1 peptidase activities. Here we report that TMK1 is cleaved by DA1 family peptidases to form a C-terminal kinase domain (TMK1C) that is relocated to the nucleus. Mutations in the DA1 cleavage site of TMK1 did not function in apical hook formation, apical hook formation was reduced in DA1 family mutants, and auxin promoted DA1 family cleavage of TMK1. This TMK1 cleavage mechanism establishes apical hook formation by a gradient distribution of auxin that controls DA1 peptidase activity, TMK1 cleavage and differential cell elongation.

## Results

### TMK1 is cleaved by DA1 family peptidases

The N-terminus of cleaved TMK1 was predicted from mass-spectrometric analysis of tryptic peptides to be between amino acid (aa) 500 - 550 (25). Examination of the TMK1 sequence revealed amino acids related to those in the DA1 cleavage site in BB (NAYK) (27), from aa 520 to aa 537 (<u>NAV</u>VVHPRHSGSDNESV<u>K</u>) (Fig. 1*A*). We therefore assessed whether DA1, DAR1 and DAR2 could cleave TMK1 near these sequences. Fig. 1*B* shows that DA1 and DAR2, and, to a lesser extent DAR1, all cleaved TMK1-3FLAG to form a C-terminal -3FLAG protein of the expected approximate 50 kDa size (25) when transiently expressed together in Arabidopsis *da1dar1* mesophyll protoplasts. Collectively this is referred to as DA1-family mediated cleavage. As DAR2 was the most active peptidase in these conditions it was used in subsequent cleavage assays. Comparison of the sizes of the DAR2 cleavage band with a TMK1-3FLAG fusion protein truncated to aa 522, the location of the predicted DA1 family cleavage site at NAVV, based on similarity to the NAYK site in BB, showed they had similar mobilities on SDS-PAGE (Fig S*1*) and was therefore close to the DA1 cleavage site. A deletion removing aa 506-537, TMK1(Δ YK) that encompassed the predicted DA1 cleavage region, was not cleaved (Figs. 1*A*, 1*C*), and a shorter deletion removing aa 521-536 (TMK1 Δ AV) was also not cleaved (Figs. 1*A*, 1*C*), defining this 15 aa region as essential for cleavage. To test this predicted cleavage region, the BB cleavage site (NAYK) and a mutated version (NGGK) that abolished DA1-mediated cleavage of BB (27) were added into the Δ AV deletion (Fig. 1A). DAR2-mediated cleavage was not observed in the NGGK sequence, while low levels of cleavage product were observed in TMK1 with the swapped NAYK cleavage site (Fig. 1*C*). This established that DA1 family peptidases catalysed the cleavage of TMK1 at a site starting at NA (aa 520-521) and extending to aa 537, consistent with the region initially identified using mass spectrometry (25), and released a similar sized 50 kDa C-terminal fragment of TMK1.

**Figure 1.**
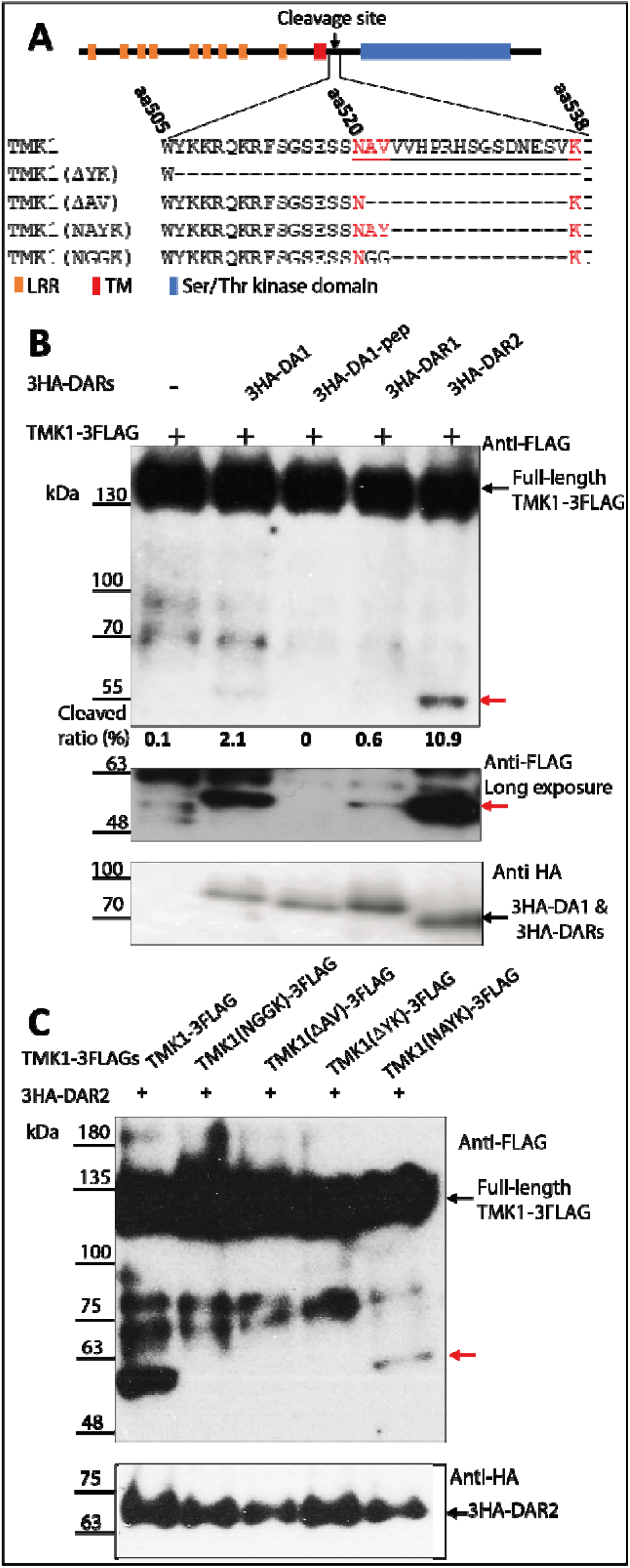
TMK1 is cleaved by DA1 family peptidases. **A**. Diagram and amino acid sequences of a cytoplasmic region of TMK1 encoding the intracellular kinase domain (blue box) adjacent to the transmembrane domain (red box). The extracellular region with Leucine Rich Repeats (LRR) are shown as orange boxes. A putative DA1 cleavage site from amino acids 520-537 is underlined. Dashes represent deleted amino acids in the different mutant TMK1 proteins used to assess cleavage. Red text represents the amino acids of the predicted cleavage site and the swapped BB cleavage site. **B**. Immunoblot of transiently expressed TMK1-3FLAG proteins together with 3HA-DA1 family members in Arabidopsis *da1-1 dar1* mesophyll protoplasts. 3HA-DA1pep is a mutant in the peptidase active site. An approximate 50 kDa TMK1-3FLAG cleaved product was observed (red arrow). A longer exposure shows DAR1-mediated cleavage. The lower panel shows 3HA-DA1, -DAR1 and - DAR2 expression levels. Cleavage levels are the % total FLAG protein in the cleaved band, shown by a red arrow. **C**. Immunoblot of transiently expressed TMK1-3FLAG mutant proteins described in panel **A** together with 3HA-DAR2 in Arabidopsis mesophyll protoplasts. Deletions of the region containing the putative DA1 cleavage site abolished cleavage, while adding the BB cleavage site to the deleted TMK1 protein recovered cleavage (red arrow). The lower panel shows 3HA-DAR2 expression levels.

### Cleavage by DA1 family peptidase facilitates TMK1C relocation to the nucleus

After cleavage, the C-terminal kinase domain of TMK1 accumulates in the nucleus and phosphorylates the auxin regulated transcriptional repressors IAA32 and IAA34 to modulate gene expression (25). To test the function of DA1 family member peptidase activities in nuclear localisation, TMK1-eGFP and cleavage deficient mutant versions were co-expressed with 3HA-DAR2 in *dar2-1* loss-of-function root protoplasts. These were used to avoid high levels of chlorophyll fluorescence. eGFP fluorescence was measured in images of protoplasts to calculate the proportion of TMK1-eGFP at the plasma membrane and in the nucleus. HISTONE H2B (HTB2) fused to mCherry (HTB2-mC) was used as a nuclear marker. TMK1C-eGFP, from aa 538 adjacent to the putative DA1 cleavage region, was used to test the TMK1C-eGFP subcellular location of the DA1 family cleavage product. Over 90% of wild type TMK1-eGFP was located on the plasma membrane when expressed without DAR2, while nearly half of the TMK1C-eGFP construct was detected in the nucleus (Figs. 2*A, 2C, 2I*). This demonstrated the expected pattern of subcellular locations of full-length TMK1-eGFP and TMK1C-eGFP in the protoplast system. Strong GFP fluorescence was detected in the nucleus when TMK1-eGFP was co-expressed with DAR2 (Figs. 2*B*, 2*I*), indicating the dependence of TMK1-eGFP nuclear location on DAR2. Three non-cleaved forms of TMK1 (TMK1(NGGK)-eGFP, TMK1(Δ AV)-eGFP and TMK1(Δ YK)-eGFP) were co-expressed with DAR2 and showed similar levels of plasma-membrane location to TMK1-eGFP in the absence of DAR2. (Figs. *2A, 2E, 2F, 2G, 2I*). TMK1-eGFP containing the swapped BB cleavage site (NAYK) when co-expressed with DAR2 accumulated lower levels of GFP fluorescence in the nucleus compared to wild-type TMK1-eGFP (Figs. 2*D*, 2*H*, 2*I*), and more than the non-cleaved TMK1-eGFP mutant proteins. These levels of nuclear TMK1-eGFP fluorescence were consistent with their cleavage efficiencies assessed using immunoblots (Figs. *1*, *2*), supporting a role for DA1-family catalysed cleavage in the nuclear localization of TMK1 C-terminal kinase domain.

**Figure 2.**
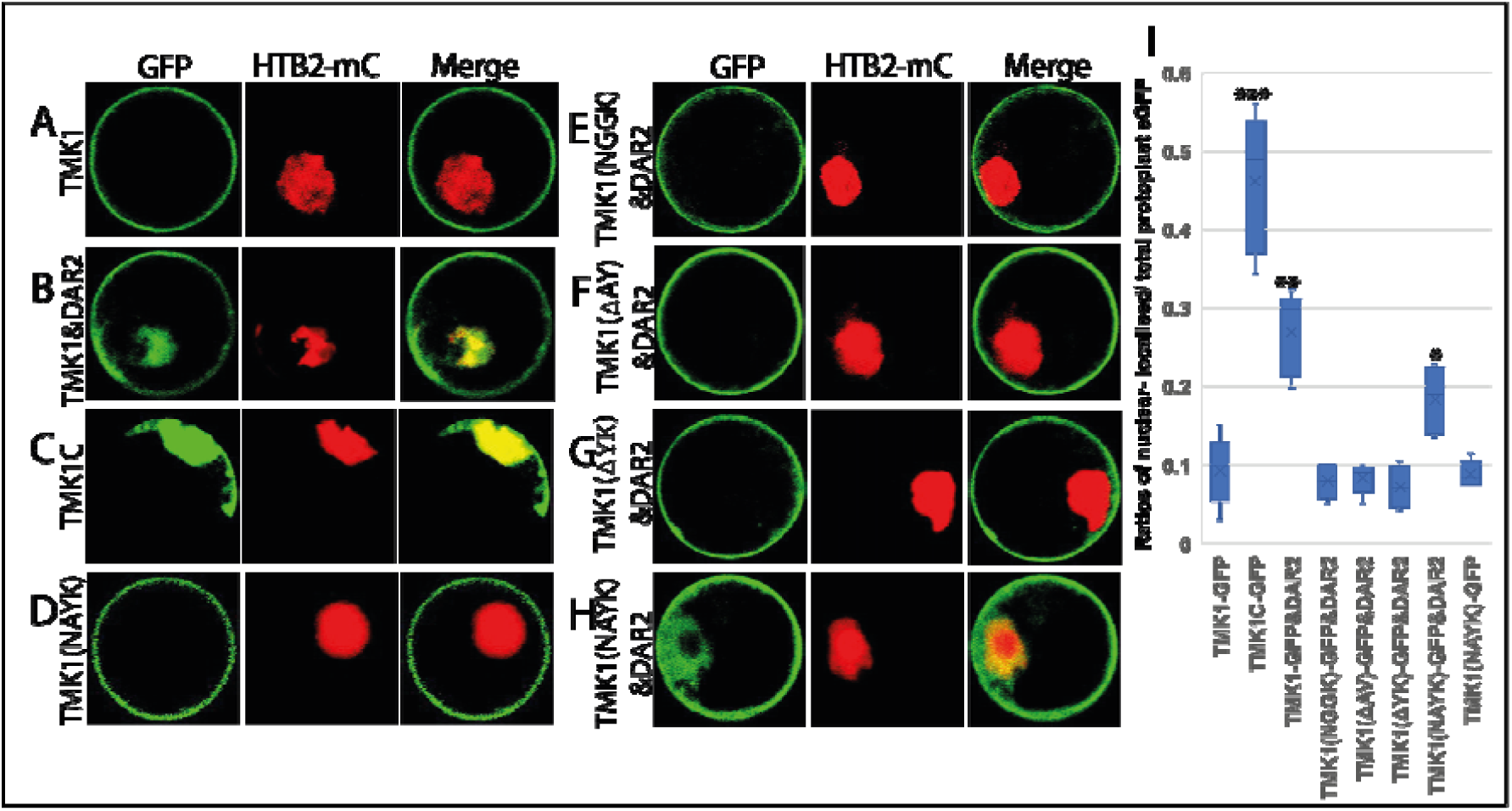
DAR2 cleavage of TMK1-eGFP relocates it from the plasma membrane to the nucleus of Arabidopsis root protoplasts. **A-H**. Confocal images of *dar2-1* root protoplasts transformed with TMK1-eGFP, Histone HT2B-mCherry and cleavage mutants with or without DAR2. HT2B-mCherry marks the nucleus. Co-location of TMK1-eGFP and HT2B-mCherry shows as yellow fluorescence. **I**. Ratios of nuclear-localised/ total protoplast eGFP in different TMK1-eGFP cleavage mutants. (*) p<0.05, (**) p<0.01, (***) p<0.001. n= 20 protoplasts from each of three independent experiments. All were compared to TMK-eGFP levels without 3HA-DAR2.

### DA1-mediated TMK1 cleavage is essential for apical hook formation

The activities of TMK1 cleavage mutants in apical hook formation were studied in transgenic *tmk1-1* lines expressing TMK1 cleavage mutants from a 2.7kb *TMK1* promoter. The *tmk1-1* mutant had an open apical hook, while wild type Col-0 seedlings had fully closed apical hooks (25). *tmk1-1* transformed by the wild type *TMK1* construct *pTMK1::TMK1* fully restored the open mutant apical hook to a closed hook indistinguishable from wild type Col-0 (Figs. 3*A*, 3*B*). In contrast, transgenics expressing the three non-cleaved forms of TMK1 (*pTMK1::TMK1(NGGK);tmk1-1, pTMK1::TMK1(*Δ *AY);tmk1-1* and *pTMK1::TMK1(*Δ *YK);tmk1-1*) displayed open apical hooks similar to *tmk1-1*, indicating a loss of function (Figs. 3*A*, 3*B*). Transformants expressing the swapped BB cleavage site construct *pTMK1::TMK1(NAYK)* exhibited an intermediate apical hook angle (Figs. 3*A*, 3*B*) consistent with reduced cleavage and/or stability of TMK1(NAYK) in protoplasts shown by immunoblots and transfected protoplast images (Figs. 1*B*, 1*C*, 2*B*, 2*H*, 2*I*). These phenotypes establish an essential role for DA1-family mediated TMK1 cleavage in apical hook formation.

**Figure 3.**
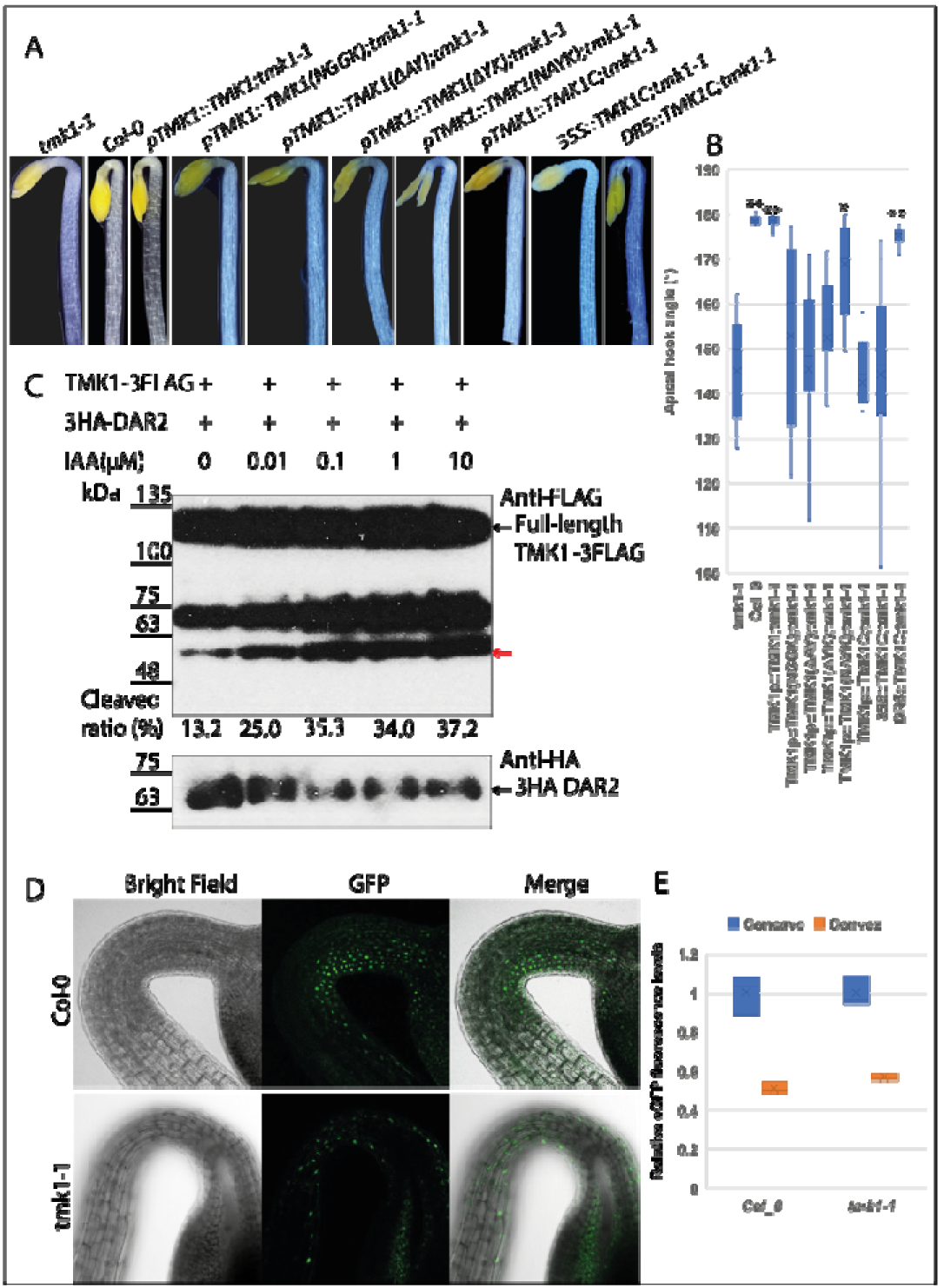
TMK1 cleavage by DA1 family peptidases is induced by auxin to form the apical hook. A. Complementation of *tmk1-1* apical hook angles by TMK1 cleavage mutants. B. Quantitative analyses of apical hook angles in *tmk1-1* transformants. n = 25 apical hooks of five independent single locus transformants were measured. (*) p<0.01, (**) p<0.001. All measurements were compared to *tmk1-1*. C. Immunoblot of TMK1-3FLAG and 3HA-DAR2 transfected *dar2* mesophyll protoplasts treated with different levels of IAA. The red arrow indicates cleaved TMK1-FLAG protein. The lower panel shows 3HA-DAR2 expression levels. D. *DR5::NLS-2xeGFP* expression levels report auxin distribution across the apical hook of Col-0 and the *tmk1-1* mutant. E. Quantitative analyses of eGFP fluorescence on the concave and convex regions of the apical hook in Col-0 and the *tmk1-1* mutant. n= 15 hypocotyls. Fluorescence levels were compared to those of the concave region of Col-0.

### An auxin-mediated gradient of TMK1C drives asymmetric apical hook growth

Apical hooks are formed by asymmetric cell elongation established by a gradient of auxin distribution, from higher levels at the concave surface to lower levels at the convex surface that drives differential cell growth (30). Nuclear-localised TMK1 is present at higher levels on the concave side of the apical hook, suggesting higher auxin levels on the concave side increase the cleavage and nuclear localization of TMK1 (25). Therefore the auxin responsiveness of DAR2-mediated cleavage of TMK1-3FLAG were analysed. 3HA-DAR2 and TMK1-3FLAG were co-expressed in *dar2* mesophyll protoplasts treated with IAA. Immunoblots showed levels of cleaved TMK1-3FLAG were progressively increased in response to higher levels of IAA in the presence of 3HA-DAR2 (Fig. 3*C*), confirming that TMK1 cleavage by DAR2 is promoted by IAA. Expression of a gradient of TMK1C by an auxin-responsive promoter such as *DR5* (31) should therefore bypass the need for DA1 family-mediated peptidase cleavage and establish normal apical hook formation. Comparison of eGFP fluorescence densities between concave and convex halves of the apical hook region of *tmk1-1* and wt Col-0 expressing *DR5::NLS-2eGFP* showed a similar distribution of nuclear-localised eGFP in both *tmk1-1* and Col-0 with higher levels seen on the concave side (Figs. 3*D*, 3*E*), indicating a negligible influence of *TMK1* on auxin distribution. Figure S2 panel A shows that expression of DA1, DAR1 and DAR2 are slightly reduced by auxin treatment in young seedlings, ruling out a role for auxin gradients influencing DA1 family -mediated cleavage by increasing their expression. *TMK1C* expression from the *DR5* promoter in the *tmk1-1* mutant formed a nearly closed apical hook, demonstrating complementation of the *tmk1-1* phenotype. In contrast, expression of *TMKIC* from the *TMK1* and the CaMV35S promoters failed to complement the *tmk1-1* open hook phenotype (Figs. 3*A*, 3*B*). Therefore, a gradient of TMK1C generated in response to auxin and DA1-family mediated cleavage is required for apical hook formation.

### *DAR1* plays major role in apical hook formation

The roles of DA1 family members in apical hook growth patterns were determined by measuring apical hook angles of single and double mutant combinations of *DA1, DAR1* and *DAR2* T-DNA loss-of-function mutants. All mutants showed varying degrees of open apical hooks (Figs. 4*A*, 4*B*), consistent with the redundant cleavage of TMK1 by these peptidases (Fig. 1*A*). Surprisingly, *dar1-1* had the most open apical hook compared to *da1-ko1* and *dar2-1* single mutants (Figs. 4*A*, 4*B*), despite 3HA-DAR1 showing the lowest TMK1 cleavage levels in protoplast assays (Fig. 1*B*). Double mutants of *da1-ko1dar1-1* and *dar1-1dar2-1* had a similar open apical hook to the single mutant of *dar1-1*, confirming the minor contributions of *DA1* and *DAR2* to apical hook formation *in vivo* (Figs. 4*A*, 4*B*). Comparison of *DA1, DAR1* and *DAR2* expression levels in dark-grown germinating seeds showed that *DAR1* expression was very high at early stages of seedling growth, while *DA1* and *DAR2* were expressed at relatively low levels (Figs. 4*C*, *S2B*). These expression patterns suggested that *DAR1* has a major role in apical hook formation.

**Figure 4.**
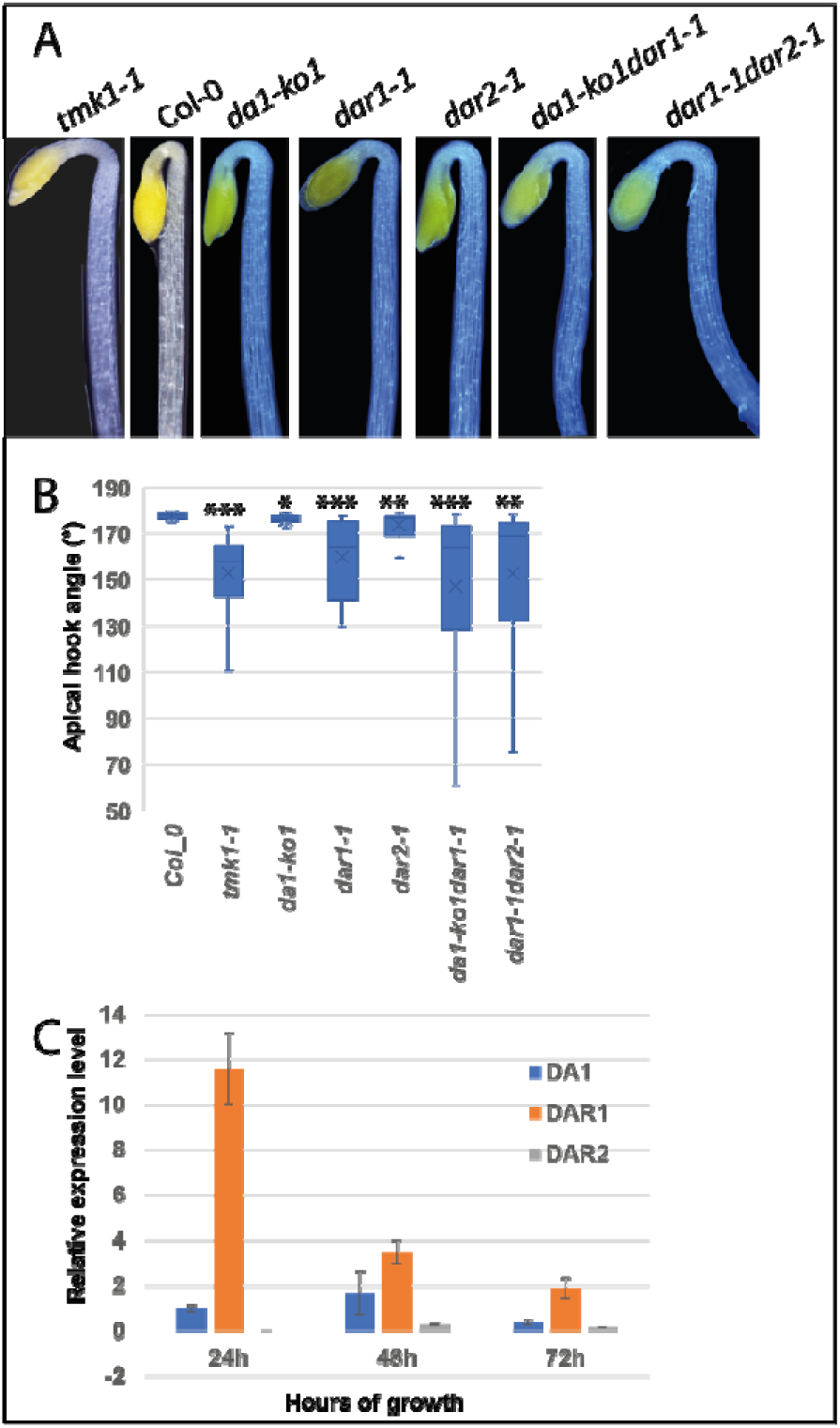
Apical hook angle of DA1 family loss-of-function mutants. **A**. Apical hook angles of *DA1* family peptidases mutants. **B**. Analysis of apical hook angles of *DA1* family peptidases mutants compared with *tmk1-1*. (*) p<0.05, (**) p<0.001.(***) p<0.0001. n= 15 hypocotyls **C**. Q-RT-PCR analyses of expression levels of *DA1* family peptidases during seed germination. Expression levels were normalised to *DA1* expression levels at 24 hrs growth in the dark.

## Discussion

Proteases and peptidases are involved in diverse processes in plants and animals, including protein turnover, senescence, cell death, defence responses and development (32). Despite these central roles, our knowledge of their regulation and substrates remains limited, particularly in plants. Here we show that members of the DA1 family of peptidases modulate the auxin-dependent cleavage and nuclear localisation of the C-terminal kinase domain of the plasma membrane localised RLK TMK1 in *Arabidopsis thaliana*. This cleavage toggles TMK1 kinase activities between its membrane and nuclear locations (33): at the plasma membrane it phosphorylates and activates H^+^ -ATPases that acidify and loosen cell walls to promote cell growth (20, 21) and initiates a kinase cascade involved in lateral root cell proliferation (23); in its nuclear location it phosphorylates and stabilises the transcriptional repressors IAA32 and IAA34, increasing repression of gene expression and limited cell growth (25, 34).

DA1, DAR1 and DAR2 cleaved TMK1 at a site that is consistent with that predicted from mass-spectrometric analyses of cleaved TMK1 *in vivo* (Fig. 1, Fig. S*1*) (25). This predicted cleavage site is similar to that of BB, which was defined by N-terminal amino acid sequencing to be between AY to form a neo-N terminal YK sequence that is targeted by the N-degron system *via* the bulky aromatic Y residue (27). The predicted TMK1 cleavage sequence has V at this position that is not targeted by the N-degron, suggesting DA1-family mediated cleavage may not directly destabilise TMK1C. Deletion of amino acids 521-536 containing the N<u>A</u>VV—<u>V</u>K sequence of TMK1 (Δ AV) abolished DA1-family catalysed cleavage, while reforming the BB cleavage site NAYK in Δ AV recovered cleavage, although to a lower level (Figs. 1*C*, 2*B*, 2*H*). This could be due to either low cleavage efficiency and/or the neo-N terminal amino acid Y that can destabilise cleaved proteins (35).

DA1 and DAR2 cleaved TMK1-3FLAG efficiently in Arabidopsis mesophyll protoplast assays, while DAR1 was less active (Fig. 1*B*). Expression patterns of these genes during very early stages of seedling formation showed that *DAR1* was very highly expressed compared to *DA1* and *DAR2* (Figs. *4C*, *S2B* and (36)), and the *dar1-1* loss of function T-DNA mutant had the largest reduction in apical hook formation compared to single loss-of-function mutants of *DA1* and *DAR2* (Fig. 4). Therefore, DAR1 probably has a predominant role in cleaving TMK1 during apical hook growth. DA1-GFP fusion proteins are located at the plasma membrane and in the cytoplasm and co-purify with plasma-membrane ATPase (29), consistent with cleaving plasma membrane-bound TMK1 on the cytoplasmic side. DA1-family peptidase activity appears to be distinct from those of conserved Regulated Intramembrane Proteolysis (RIP) proteases (37) such as gamma secretases (4), which cleave within trans-membrane domains (5, 29).

Peptidases and proteases catalyse irreversible reactions and their activities remain latent until needed. For example, papain-like proteases are produced as auto-inhibited pre-proteases that are activated by cleavage (32). DA1, DAR1 and DAR2 peptidase activities are controlled by mono-ubiquitylation and de-ubiquitylation (27, 28) mediated by Ubiquitin Interaction Motifs (UIM) that characterise DA1, DAR1 and DAR2 family members. DA1 peptidase activity is inhibited by phosphorylation of conserved serine and threonine residues close to its active site by the RLKs BRI1 and BAK1 (29). During apical hook growth the BR biosynthesis gene *CPD* is expressed more highly on the convex surface (38), where TMK1 cleavage is low. Conceivably, BR inhibition of DA1 family peptidase activities on the convex surface may contribute to the gradient of TMK1 cleavage. Additionally, brassinolide (BL) treatment induces the phosphorylation of the same T947 residue of the plasma membrane H+-ATPase as TMK1 to promote hypocotyl cell elongation by the acid-growth pathway (39), consistent with BR inhibition of DA1 peptidase activity and the maintenance of membrane-associated TMK1 kinase activities. Thus, DA1 family peptidase activity may form a link between auxin and BR in cell growth control.

Apical hook formation depends on the asymmetric distribution of efflux carriers in hypocotyl cells creating an auxin gradient across the emerging hypocotyl (40, 41). Lower auxin levels increase cell expansion on the convex surface of the hook and higher auxin levels decrease cell expansion on the concave surface to establish and maintain the hook angle. *TMK1* is necessary for hook formation, increased auxin levels promote cleavage to form TMK1C, and levels of nuclear-localised TMK1C increase on the concave surface where it reduces cell growth (25). DAR2-mediated cleavage of TMK1 was increased by auxin treatment (Fig. 3*C*) and expression of TMK1C from the *DR5* auxin-responsive promoter nearly fully complemented the hook angle in the *tmk1-1* loss of function mutant (Figs. 3*A*, 3*B*). In contrast, expression of TMK1C from the *TMK1* or 35S promoter did not rescue normal *tmk1-1* hook angles (Figs. 3*A*, 3*B*), showing that the graded distribution of TMK1C from low levels on the convex surface to high levels on the concave surface drives differential cell elongation and hook formation. As the expression of *DA1* family members is not increased by auxin treatment (Fig. *S2B*), this supports the idea that auxin levels modulate DAR1 peptidase activity across the hook region to generate the gradient of TMK1C levels. How auxin is sensed in this system is not yet clear, while a role for TMK1 in direct auxin perception remains unresolved (22, 24, 42).

DA1 and DAR1 peptidase activities limit the duration of cell proliferation during early stages of organ growth by cleaving and destabilising a variety of proteins that promote cell proliferation (26, 27). Initial analyses of loss-of-function mutants of *tmk1* and *tmk4* showed they act redundantly to reduce organ size by limiting cell expansion (43). TMK1 also promotes auxin-mediated leaf pavement cell growth by activating ROPs (Rho-like guanosine triphosphatases) at the plasma membrane (22). As TIR/ABF auxin signalling pathways have a central role in cell specification and organ growth (44), it is possible that DA1-family peptidase cleavage of TMK1 could influence auxin responses more generally during plant growth.

## Materials and Methods

### Plant materials

*Arabidopsis thaliana (L*.*) Heynh*. Columbia-0 (Col-0) was the wild-type line used. Mutants used were *tmk1-1*(SALK_016360), *da1-ko1*(SALK_126092), *dar1-1*(SALK_067100), *dar2-1*(SALK_016122), *da1-ko1dar1-1* and *dar1-1dar2-1* (26). Transgenic plants were made in the *tmk1-1* background. *DR5::NLS-2xeGFP* transgenics were made in Col-0, and crossed into *tmk1-1* to generate *DR5::NLS-2xeGFP tmk1-1*.

### Plant growth

Plants were grown in controlled growth cabinets at 20°C under long day conditions of 16h light/8h dark.

### Apical hook measurements

Sterilised Arabidopsis seeds were sown on ½ MS (Murashige and Skoog) medium without glucose, kept at 4°C in the dark for 4 days, then grown vertically at 20°C in the dark for 60-65 hr. Images were taken with a ZEISS Axio Zoom V16. Apical hook angles were measured using Image J in 15 individuals for each line.

### Protoplast extraction and transformation

Leaf mesophyll protoplasts were made from 3-4 week - old *da1-ko1dar1-1* or *dar2-1* plants grown under 8h light/16h dark at 20^0^C and prepared and transfected according to (45). Root protoplasts were made from *dar2-1* seedlings grown on vertical MS agar plates for 10-14 days. Roots were excised and finely chopped in 20 mM MES pH5.7, 20 mM KCl, 10 mM CaCl_2_, 0.4 M mannitol, 2.5% cellulase R10, 0.65% macerozyme (Yakult), degassed three times for 1 min, then digested with gentle agitation at 20°C for 14h. Protoplasts were filtered through 70 μm then 40 μm mesh and collected by centrifugation at 250g for 6 mins. Protoplasts were transfected with PEG according to (45). Leaf protoplasts were cultured in W5 medium for 16h and total protein was extracted for immunoblotting. Transfected root protoplasts were cultured in W5 for 30h prior to imaging. For IAA treatment, while transfected leaf protoplasts were grown in W5 medium for 16 hrs and IAA was added 3 hrs before extraction.

### Imaging

A Leica SP8X with an HC Plan Apo 20x/NA 0.75 water immersion objective lens was used for confocal imaging. eGFP was excited at 488 nm with an Argon ion laser and 500 nm to 550 nm emission was collected. mCherry was excited at 561 nm with a white-light laser and 590 nm to 650 nm emission was collected. The confocal pinhole size was 1AU at 525 nm. Confocal images of TMK1-eGFP distribution in protoplasts were analysed by defining nuclear areas by the HTB2-mCherry signal and the whole cell area was defined by the TMK1-eGFP signal. eGFP fluorescence was measured in the nuclear and whole cell areas by Image J. The TMK1-eGFP cleavage ratio was calculated as the nuclear eGFP signal / eGFP signal in the whole cell. Five protoplasts were measured in each of three independent transformations.

### Cleavage ratio measurement

Immunoblots of cleaved TMK1 were measured using Image J. The cleavage ratio was calculated as the intensity of the cleaved protein / intensities of the cleaved and full-length protein.

### Auxin distribution in apical hook

GFP fluorescence in apical hooks was measured in concave and convex areas (divided by the central vascular bundle) using Image J. Three individuals were measured for each genotype.

### Gene accessions

The genes used in analyses were *TMK1* (AT1G66150), *DA1*(AT1G19270), *DAR1*(AT4G36860), *DAR2* (AT2G39830), *BB* (AT3G63530) and *HTB2* (AT5G22880).

### Plasmid construction

Plasmids *35S:3HA-DA1, 35S::3HA-DA1-pep, 35S::3HA-DAR1* and *35S::3HA-DAR2* in pB7GW2 were described in (27). To create expression vectors for protoplast transfection, the coding region and intron of TMK1 was amplified from Col 0 seedling genomic DNA, inserted into pENTR/D-TOPO (ThermoFisher K240020), and cloned into the expression plasmids pW1211 and pS103 by LR reactions to generate *35S::TMK1-3FLAG* and *35S::TMK1-GFP*. pS103 was modified from pEarly103 by replacing mGFP5 with mEGFP. Mutants were generated by paired primers (Supplementary Table 1) to amplify TMK1 in pENTR/D-TOPO, then ligated by Infusion reactions (Takara 638909). A 2.7 kb region upstream of the TMK1 initiator ATG was inserted into TMK1-pENTR/D-TOPO by Infusion reaction and transferred into vector pW0211, which was modified from pw1211 by removing the 35S promoter to generate *pTMK1::TMK1-3FLAG*. The *DR5* promoter was inserted into TMK1C in pENTR/D-TOPO by infusion, then transferred into pW0211 to generate *DR5::TMK1C-3FLAG*. The *HTB2* coding region was amplified from Col_0 seedling cDNA using primers (Supplementary Table 1) and inserted into a level 0 module and *35S::HTB2-mCherry* was assembled using Golden Gate methods into the pAGM4723 ccdB vector (46).

### RNA extraction and quantitative RT-PCR

Seedlings were ground to a powder in liquid nitrogen and RNA was purified with an RNeasy Mini Kit (Qiagen 74104). Reverse transcription was performed by Superscript IV Reverse Transcriptase (Life Technologies Ltd 18090050) with Oligo(DT)15 Primer (Promega C1101). Q-RT-PCR reactions used Lightcycler 480 SYBR Green I Master Mix (Roche 04707516001) on a Roche Lightcycler 480. *Actin2*(*AT3G18780*) was used as the internal control. The calculation used the method of Zhao and Fernald (47).

### Statistical Analyses

Student’s t-test was used to compare samples with controls. Significance levels were set at p=0.05, p=0.01, p=0.001 and p=0.0001. Graphs were plotted using Excel. Error bars are SE.

## Data Availability

All data and resources described are included in the article or S1 Appendix. All constructs and know-how are available from the authors.

## Acknowledgements

We thank Dr Dolf Weijers (Wageningen University) for providing the *DR5::NLS-2xeGFP* transgenic line and Dr Eva Wegel for advice on confocal imaging. This work was funded by the Biological and Biotechnological Sciences Research Council (BBSRC) Newton Fund to BG and MWB, BBSRC Grant BB/K017225 to MWB, and Institute Strategic Grant GEN (BB/P013511/1) to MWB.

## Author Contributions

BG and MWB developed the research programme, BG HD and CS conducted experiments and analyses, and BG and MWB wrote the paper with input from all authors.

## S1 Appendix

**Figure S1. Relative sizes of the cleaved TMK1 protein and TMK1 truncated from aa 522.**

**Figure S2. Expression levels of DA1, DAR1 and DAR2 in response to auxin and during seed germination.**

**Table S1. Primers Used**

**Table S2. Q-PCR primer amplification efficiencies.**

