## Supplemental Information for "Modulation of Receptor-like Trans-Membrane Kinase 1 nuclear localisation by DA1 peptidases in Arabidopsis"

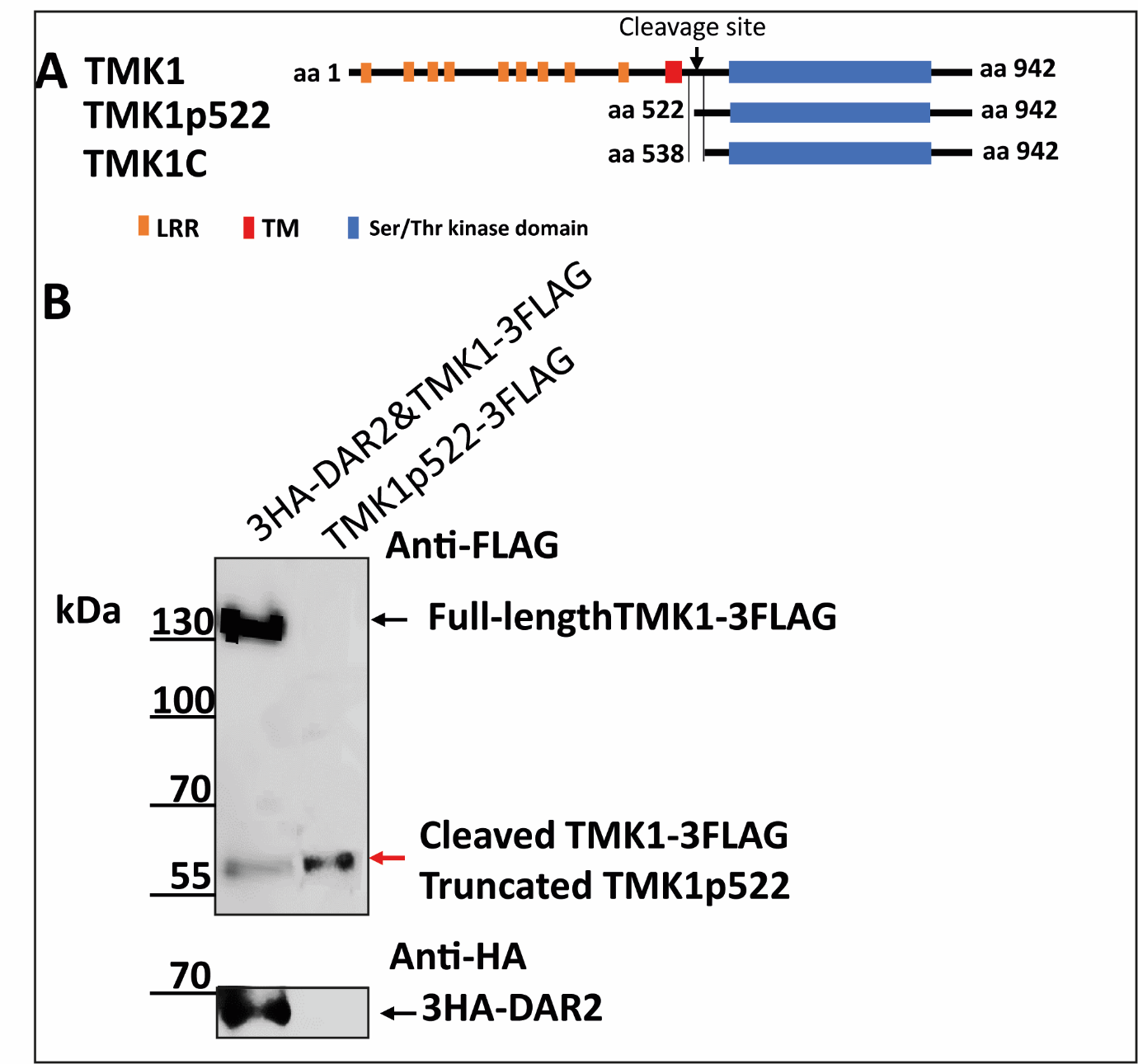

**Figure S1. Relative sizes of the cleaved TMK1 protein and TMK1 truncated from aa 522.**

A. The structures of TMK1, TMK1p522 (truncated from aa 522) and TMK1C. The Intracellular kinase domain (blue box) is adjacent to the transmembrane domain (red box). The Leucine Rich Repeats (LRR) (orange boxes). The start and end amino acid (aa) of each protein is indicated.
B. Immunoblot showing the relative sizes of cleaved TMK1-3FLAG and truncated TMK1p522-3FLAG.

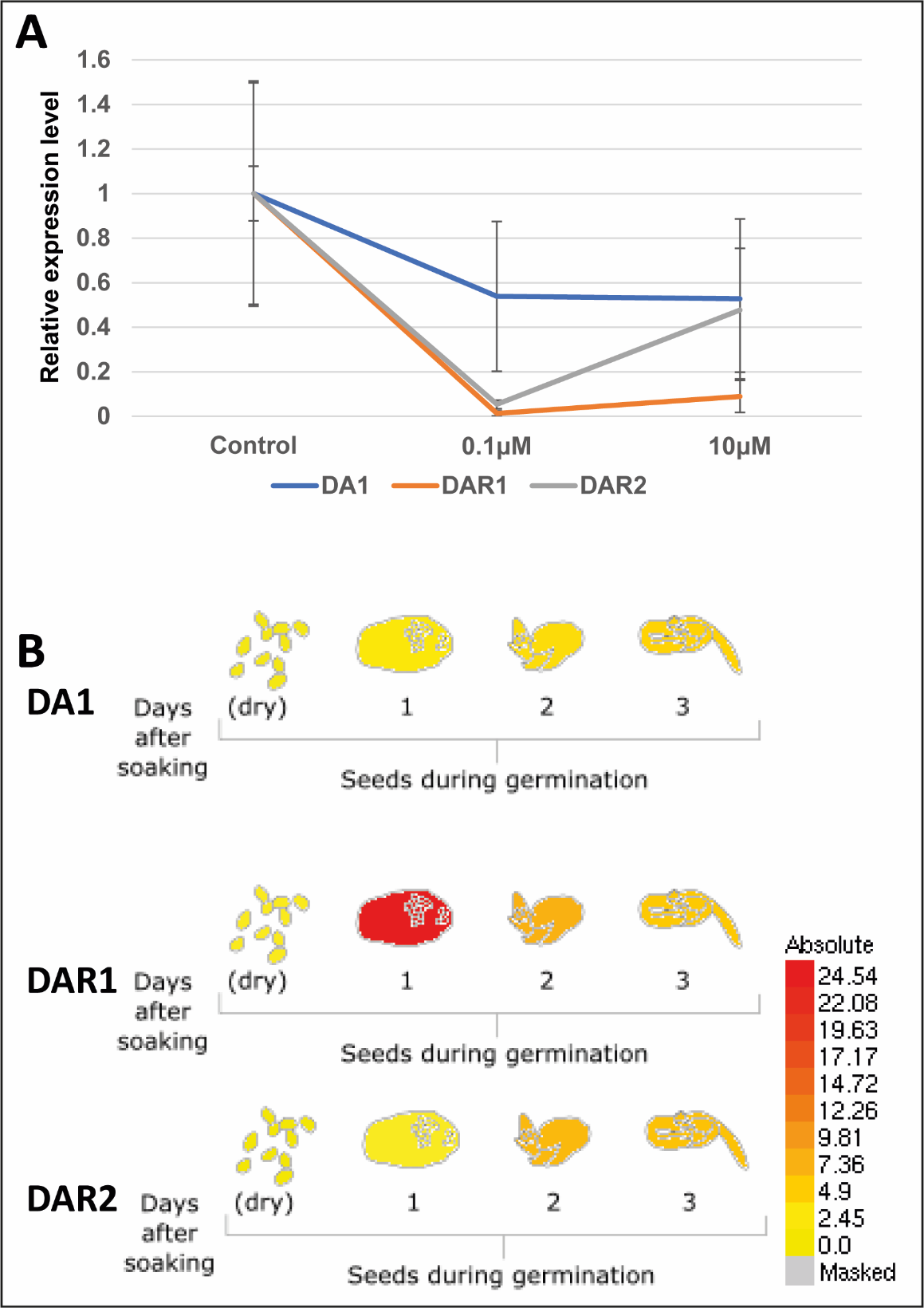

**Figure S2. Expression levels of DA1, DAR1 and DAR2 in response to auxin and during seed germination.**

A. *DA1* family peptidase expression levels are not induced by auxin. Seedlings were treated with 0.1μM or 10 μM IAA for 4 hrs, and ethanol was used as a control. Q-RT-PCR data were normalised to control treated seedlings.
B. Expression pattern of *DA1* family peptidases during seed germination. Image from https://www.arabidopsis.org/.

| **Table S1. Primers** |  |  |
| --- | --- | --- |
| pENTR/D-TOPO | Direction | Sequence |
| TMK1 | Forward | CACCATGAAGAAAAGAAGAACCTTTCTTCTAT |
| TMK1p522 | Forward | CACCATGGTAGTGGTGCATCCGC |
| TMK1C | Forward | CACCATGATTACAGTTGCGGGTTCAAG |
| TMK1 | Reverse | TCGTCCATCTACTGAAGTGAATG |
| In-fusion | Direction | Sequence |
| TMK1p1 | Forward | ATGAAGAAAAGAAGAACCTTTCTTCTAT |
| TMK1p538 | Forward | ATGATTACAGTTGCGGGTTCAAG |
| pENTR/D-TOPO-R | Reverse | GGTGAAGGGGGCGGCCG |
| TMK1promoter | Forward | GCCGCCCCCTTCACCCTTTCTGATCCTTATTTTTTGCT |
| TMK1promoter | Reverse | TCTTCTTTTCTTCATAGCTTGAAGAAACAGAGG |
| TMK1promoter_TMK1C | Reverse | CGCAACTGTAATCATAGCTTGAAGAAACAGAGG |
| TMK1(ΔYK) | Forward | CTGTTGGATTACAGTTGCGGGTTC |
| TMK1(ΔYK) | Reverse | ACTGTAATCCAACAGAAAACTAACAAACC |
| TMK1(ΔAV) | Forward | GCTCAAATAAGATTACAGTTGCGGG |
| TMK1(ΔAV) | Reverse | TAATCTTATTTGAGCTCTCACTTCC |
| TMK1(NAYK) | Forward | AATGCATACAAGATTACAGTTGCGGG |
| TMK1(NAYK) | Reverse | AATCTTGTATGCATTTGAGCTCTCAC |
| TMK1(AV-GG) | Forward | CAAATGGAGGAAAGATTACAGTTGCGGG |
| TMK1(AV-GG) | Reverse | TCTTTCCTCCATTTGAGCTCTCACTTCC |
| DR5 | Forward | GCCGCCCCCTTCACCGAATTCGTCGACGGTATCGC |
| DR5_TMK1C | Reverse | CGCAACTGTAATCATTGTTATATCTCCTTGGATCGATCC |
| GoldenGate | Direction | Sequence |
| HTB2 | Forward | tgtgaagacaaAATGGCGAAGGCAGATAAGAA |
| HTB2 | Reverse | tgtgaagacaaCGAAGAACTCGTAAACTTCGTAACC |
| Quantitative RT-PCR | Direction | Sequence |
| Actin2 | Forward | GAAATCACAGCACTTGCACC |
| Actin2 | Reverse | AAGCCTTTGATCTTGAGAGC |
| DA1 | Forward | CACGGCAATGTGAAGTTACC |
| DA1 | Reverse | ATCCTTTGAGCCTCATCCAC |
| DAR1 | Forward | GTGTCAGCATCGTCCAAGAA |
| DAR1 | Reverse | TCGCCTCAGACCATGCTTAA |
| DAR2 | Forward | CTTCAACATCATCGGTGGCC |
| DAR2 | Reverse | CATTTGCTGCCCTGAAACCT |

| **Table S2. Q-PCR primer amplification efficiencies** | |
| --- | --- |
| Gene Name | Amplification Efficiency |
| ACTIN2 | 2.0823 |
| DA1 | 2.0160 |
| DAR1 | 1.9617 |
| DAR2 | 2.3064 |
